# Bridging Bibliometrics and Biology: A Framework for the Relevance Scoring and Biological Stratification of Scalable MeSH Co-occurrence Networks

**DOI:** 10.64898/2026.10.29.762716

**Authors:** Jakob Sax, Linda Schenk

**Affiliations:** Institute of Environmental Medicine (IMM), Karolinska Institutet, Stockholm, Sweden

## Abstract

Systematic literature review is often limited by the rigidity of search strings, which often miss non-obvious but highly relevant topics. Controlled vocabularies such as Medical Subject Headings (MeSH) facilitate the review process, but relationships between terms, and their relevance to a hypothesis of interest, are rarely stated explicitly. Presented here is a MeSH term extraction and networking technique, seeded by a single query for Allergic Contact Dermatitis, forming a “themed” network of terms. The resulting co-occurrence network is pruned by an unsupervised consensus of Metropolis-criterion likelihood search strategies, isolating a significant core subgraph and filtering out incidental co-indexing. Symmetrically normalized Article Relevancy Scores (ARS) then evaluate each article against the network’s weighted terms across the annotated MEDLINE corpus, correcting for publication volume bias; Mean Relevancy Scoring (MRS) inverts these scores back onto the terms, scoring each MeSH term by the mean ARS of the articles carrying it to measure its specificity to the query. Benchmarked against the expert-curated Adverse Outcome Pathway (AOP) for Allergic Contact Dermatitis, the network recovered key event terminology and identified highly specific literature that standard retrieval methods overlooked. This data-driven approach quantitatively assesses associated MeSH terminology, extracting biologically relevant concepts and key relationships directly from the underlying structure of the literature. By automating the discovery of these latent connections, the framework provides a reproducible, scalable foundation for understanding the biomedical literary space surrounding a given search string, assisting systematic review creation, mechanistic pathway mapping, and hypothesis generation across biomedical disciplines.

## 1. Introduction

The rapid sustained growth of biomedical literature, presents a profound challenge for researchers organizing knowledge, conducting systematic reviews, or sorting complex multi-level biological processes. While, databases such as MEDLINE, a core component of PubMed^1^, play a pivotal role in systematically indexing^2^ this vast corpus, manual review remains the primary method for sifting through these databases. This manual reliance inherently limits the scope of discovery and risks omitting non-obvious, cross-disciplinary connections. Existing computational efforts attempt to mine this literature using tools such as, Named Entity Recognition (NER)^3–6^ for molecular interactions or Multi-Level Event Extraction (MLEE) to attempting to capture broader biological contexts. ^7–6^ While these natural language processing approaches successfully extract complex interactions to form large-scale knowledge graphs, applying deep semantic parsing across the entirety of MEDLINE is computationally exhaustive. Addtionally, these methods rely on static copies of ontologies and risk becoming antiquated. Furthermore, extracting millions of localized semantic events does not inherently resolve the topological relevance of those events. Approaches attempting to resolve semantic relationships within a 33-million article corpus generate massive, unpruned networks. They therefore require built-in mechanisms to dynamically score and filter individual article contributions based on global network architecture.^10^

Bridging this informational gap requires methodologies that move beyond the extraction of localized syntactic events to establish throughlines of complex structural associations within the corpus. These methodologies should prioritize accessibility, enabling researchers to rapidly generate systemic knowledge graphs from broad conceptual queries without relying on highly specialized, predefined target pairs. Such accessible and efficient contextualization is critical for hypothesis formulation. manually organizing knowledge into a simplified semi-linear structures whether through graph theory^11^, biological organizational charts, or Adverse Outcome Pathways (AOPs)^12^, is an exhaustive task. These constructed flows represent complex multi-faceted hypotheses nested within larger unrepresented networks of interactions. Identifying these latent relationships and assessing them within the context of parallel associations allows researchers to systematically connect “lower” events (e.g. molecular specific gene transcription) to “higher” level outcomes (e.g. diseases). A context-driven understanding of the literature is therefore paramount for prioritizing evidence and comprehensively mapping the cascading implications of a specific query.

To achieve this comprehensive contextualization, literature indexed under the controlled ontology of Medical Subject Headings (MeSH) provides a highly curated and continuously updated dataset with which to evaluate the ongoing state of biomedical literature. MeSH terminology evolves with the scientific literature and is continuously applied to articles using both machine learning techniques as well as human indexers^13^, providing a robust feature dataset for assessing core concepts applied to these articles. Several efforts have leveraged MeSH co-occurrence within articles to construct topical networks, utilizing probabilistic models for bibliometric analysis^14–16^ or visualization techniques to map research trends in specific domains, such as pharmacology^17,18^ and systemic disease relationships.^16^ Advanced frameworks, have also demonstrated the utility of gene to MeSH enrichment networks within specific literature subsets.^20^ While these localized approaches map specific domains or entity pairings, they generally fail to capture the dynamic bidirectional relationship between a curated conceptual network and the broader biomedical corpus from which the sub-network is derived. There remains a need for a methodology capable of contextualizing and evaluating how hierarchical concepts bibliometricly interact and influence one another, and to quantitatively score the relevance of key terms and individual articles based on their influence within the identified sub-network, while accounting for general bibliometric prevalance.^21^ By applying unbiased statistical consensus pruning to vast, over-dispersed datasets, researchers can isolate highly relevant theme-centric subgraphs while still being able to discriminate obscure yet connected topics.

Following the identification of a theme-centric subgraph, an additional computational challenge emerges in assessing the relevance of individual article semantic contributions to the subgraph structure, while maintaining context within the global dataset. We postulate this can be resolved by applying a relevance scoring metric that bridges global network topology with localized subgraph architecture. Specifically, an article’s contextual relevance can be mathematically scored based on the intersection of its constituent MeSH terms with the pruned subgraph. While initial term weights correspond to global baseline centrality, raw summation is highly susceptible to publication volume bias. To correct this, a symmetrically normalized projection explicitly dampens the advantage of heavily indexed articles. Articles aggregated under specific MeSH nodes can then be cumulatively assessed for their true topological influence while maintaining contextualization to the global dataset. This dual-layered approach enables the automated prioritization of biomedical literature based directly on an article’s topological contribution, accurately tracking a MeSH term’s specific influence rather than mere corpus ubiquity.

While network topology and mathematical scoring can drive discovery, these structural metrics do not natively convey how biological processes relate conceptually. It is therefore highly beneficial to assign strata (e.g. biological organization) labels to MeSH terms to serve as a complementary interpretive layer. This addition yields insights into which biological levels exert a heavy influence in a particular themed network.

To demonstrate the practical utility of this networking contextualization and mapping approach to biomedical literature review we demonstrate proof-of-concept through the extraction, categorization and quantification of MeSH terms generated from a pre-specified MEDLINE query. Applied to the well-understood biological process of skin sensitization^22^, this pipeline is applied to a query of the MeSH term “Dermatitis, Allergic Contact”^23^ to evaluate structural and terminological recovery objectively against an expert-curated ground truth network. Stratification is subsequently applied to the final subgraph to reflect biological levels of organization commonly found within AOPs.^24–26^ Ultimately, this approach yields an unbiased, contextualized semantic network of MeSH terms, providing structural insights into bibliographic associations to assist in identifying relevant literature for AOPs and biomedical hypothesis generation.

## 2. Methods

### 2.1. Data collection & processing

A *Medical Publishing to Network* (MPN) desktop application was developed to facilitate the retrieval of PMIDs and networking of MeSH terms in a user-friendly interface. To validate this Python-based MPN pipeline, the search string, “Dermatitis, Allergic Contact [Mesh]” AND 1650/01/01:2025/06/01[EDAT]’ (henceforth, “ACD query”) was applied. Citation data was retrieved for each initial PMID and assigned a generation label (P_0_=0, G_1_=1, … G_n_=*n*). Iterations for each subsequent generations maintained duplicate assignments to the oldest generation. The ACD query network was formed on P_0_+G_1_. PMIDs were parsed to their respective associated MeSH terms, extracting a comprehensive list from the NLM’s official MeSH repository^27^ to identify stop words, resulting in 30,656 unique MeSH terms. Model pipeline is outlined in **Figure 1**. Claude Code *Opus 5.5* (used for MPN pipeline user interface development and code cleaning and error identification and resolution.

**Figure 1.**
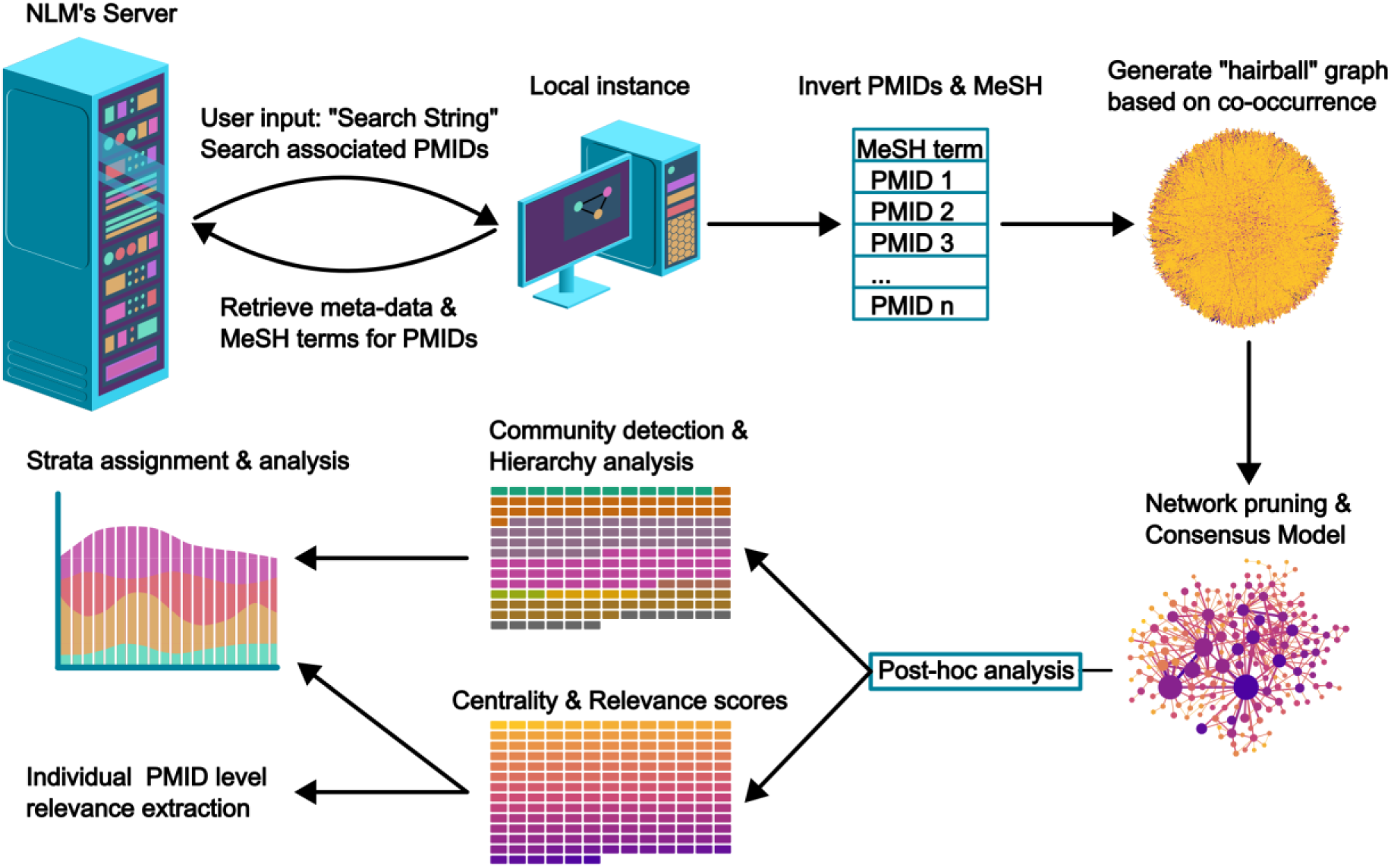
Illustration of data collection and processing for network creation from the National Library of Medicine (NLM) public database. Briefly, literature retrieval from NLM’s MEDLINE database is performed from any given query. Then the system evaluates the incoming and outgoing citations based on a user predefined generation count to evaluate further possible relevant articles. Subsequently, it categorizes all retrieved articles under their pre-associated Medical Subject Headings (MeSH) terms and generates a network based on co-occurrence (edges) of MeSH terms (nodes) within a given article. It then calculates various weights to both edges and nodes based on criteria such as normalized citation impact of articles, number of co-occurrences, centrality, etc. for the given whole network. Consensus modeling allows for the identification of the Largest Connected Component (LCC) of the most important edges. Finally, the PubMed Identifiers (PMID) for any given identified MeSH term or relationship are extracted and assigned a relevancy score with respect to its structural importance to the LCC.

#### 2.1.1. Network construction

PMIDs were mapped to associated MeSH terms using the *Bio.Entrez* package accessing the MEDLINE database (n_articles_ = 34,280,964). To ensure biological relevance only terms within the MeSH tree categories were retained for network building: ‘A’: “Anatomy”, ‘C’: “Diseases”, ‘D’: “Chemicals and Drugs” or ‘G’: “Phenomena and Processes”. External terms were categorized as stop words; removing 11,256 unique terms (**Supplementary Figure 1**). Terms were standardized by stripping “Subheading(s)” and major topic indicators to allow merging under primary MeSH terms, removing their artificial inflation of co-occurrence. Edges were defined by the co-occurrence of standardized MeSH terms within the same article.

#### 2.1.2. Zero-truncated Negative Binomial (ZTNB) regression

Co-occurrence count distributions within the unfiltered network were assessed using a Zero-Truncated Negative Binomial (ZTNB) regression model (*statsmodels* package; TruncatedLFNegativeBinomialP, truncation = 0) fitted with maximum likelihood (L-BFGS, ≤200 iterations, intercept = log of sample mean, α = 0.1) to estimate the dispersion parameter (α) (**Supplementary Table 1)**. The extracted network consists of observed edges (lower bound *Y* ≥ 1). Zero-truncation is imposed through the likelihood to estimate the latent untruncated distribution parameters from the observed positive counts without inferring unobserved structural zeros or shifting counts. Fitted over the full 1,181,631-edge population. A significant LR value indicates overdispersion relative to a Poisson process, motivating the likelihood-based filtering applied downstream. ^28^

### 2.2. Raw network pre-computations

#### 2.2.1. Weighting metrics

Node weighting metrics such as: article centrality, rank normalized article count, median citations, and total citations were calculated as percentile ranks; article-level citation counts were ranked in the same way. Unique term co-occurrences defining edge strength were log(1+x) transformed and min-max normalized. These weighting factors were calculated prior to network pruning for *post-hoc* analysis. Non-weighted metrics for nodes and edges including, the Jaccard index^26^, edge and node betweenness centrality^30,31^, Eigenvector centrality^32^, PageRank^33^, clustering coefficient^34^, and Marginal Likelihood Filter (MLF)^35^ were calculated post-network formation using the *networkx* and *community* packages.

### 2.3. Network pruning

#### 2.3.1. Stochastic Consensus Model

To identify meaningful connections between MeSH terms within the over-dispersed raw co-occurrence network from, a two-stage filtering protocol was implemented. First, significant subgraphs were independently identified using two stochastic search strategies: Global Likelihood Filter (GLF)^35^ and Simulated Annealing (SA)^36^. Total log- likelihood of the network topology was minimized independently using the Metropolis algorithm.^37,38^ Each method computed 5×10^6^ iterations, targeting an n = 500 edge count (**Supplementary Figure 2**). GLF maintained a fixed temperature (*T* = 1) while SA utilized a geometric cooling schedule of α = 0.9999995 and an initial temperature of 5000 to avoid local minima. A Consensus Model (CM) was built from the intersection of the subgraphs from these two approaches (*E_consensus_* = *E_GLF_* ∩ *E_SA_*).

#### 2.3.2. Largest Connected Component (LCC)

The resulting CM produced a fragmented graph, thus only the Largest Connected Component (LCC)^36^ was retained using *networkx* to isolate a continuous core of significant relationships for pathway analysis.. Surviving, secondary networks were examined for their size, number of nodes and edges, and “themes” to ensure target edge count predeterminants were appropriately chosen for the GLF and SA methods.

### 2.4. Post-hoc analysis

We developed a novel, literature-based weighting metric: the Article Relevance Score (ARS) and Mean Relevance Score (MRS) to holistically assess the topical relevance of each article and MeSH term against the network structure.

#### 2.4.1. Article Relevance Score

Following retrieval of all PMIDs related to MeSH terms present in the LCC, an ARS (*S_article_*) was calculated for each article by traversing the corpus twice: an initial document-frequency pass establishing term degrees, followed by a scoring pass. Let *K*denote the set of all final LCC nodes. For each node *i* ∈ *K* receives an initial weight, *c_i_*, assigned based on a predetermined intrinsic centrality metric (i.e. PageRank) from the initial unfiltered network. Each article, *A*, is annotated with its specific MeSH terms *M_A_*. ARS is the fractional sum of *c_i_* weights intersecting sets *K* and *M_A_* (*K∩M_A_*):

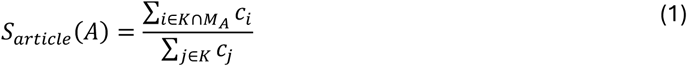

To dampen corpus-wide volumetric bias from article term counts a symmetrically degree-normalized projection was employed to reduce noise. Defining *d_A_* = |*K* ∩ *M_A_*| as the network degree of article *A*, and *d_i_* = |*P_i_* | as the degree of term *i* within the scored set (where *P_i_* is the set of articles in the corpus containing term *i*), the Normalized ARS (*S*^^^*_article_* (*A*)) is formulated as a single-pass bipartite projection:

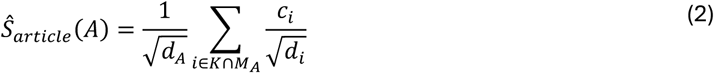

To evaluate the pathway-specific MeSH term importance, this projection is inverted back to the term level to calculate the Mean Relevancy Score (MRS). For any given network term *t* ∈ *K*, the MRS (*S̅_term_*(*t*)) represents the max-scaled, average Normalized ARS of all articles containing that term:

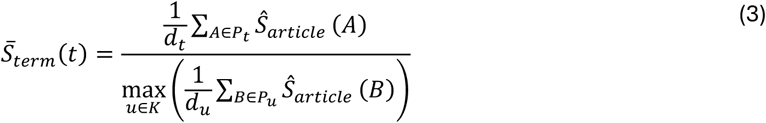

Constructing MRS as a plain mean prevents weighting advantages from high occurrence volumes, ensuring that the metric tightly tracks pathway specificity rather than ubiquity. Max-scaling bounds the distribution within [0,1], rendering the MRS invariant to underlying ARS constant scaling factors. Every *P_t_* is non-empty (*d_t_* ≥ 1), as nodes are inherently produced by the corpus.

#### 2.4.2. Ground Truth Construction and Validation

The filtered consensus network was benchmarked against a ground-truth dataset of MeSH terms derived from curated OECD Allergic Contact Dermatitis (AOP 40) reference articles^22^ to ensure appropriate identification of biological topics related to Allergic Contact Dermatitis. References were excluded if they were (i) not indexed articles (book chapters, governmental documents) or (ii) lacked MeSH indexing. Of 118 entries, this yielded 96 curated positives and 22 exclusions. Applying the stop-word filter to extracted MeSH terms yielded 256 eligible terms. A ground-truth co-occurrence network was built from terms appearing in ≥2 reference articles. Corpus prevalence was estimated from a uniform random background sample of 50,000 annotated articles (seed = 42), applying a 0.5 continuity pseudo-count for absent terms to compute base-rate-corrected enrichment. Agreement was quantified by: nodes attestation, the intersection of consensus nodes with the eligible ground-truth vocabulary; and edge recovery, the consensus edges co-occurring in ≥1 and ≥2 ground-truth articles.

#### 2.4.3. Retrievability Benchmarking and Scoring Projections

Article retrieval performance was evaluated across a defined pool (n = 8,878,867), comprising all annotated MEDLINE corpus articles containing ≥1 consensus network term (25.6% of the total corpus). Records lacking network terms score zero universally and were excluded to rank all scorers on an identical set. Of the 96 curated positives, 86 fall within this pool, establishing a retrievability ceiling of 86.6% that bounds all methods. The restriction is conservative; recomputing Boltzmann-Enhanced Discrimination of Receiver Operating Characteristic (BEDROC) over the unfiltered corpus raises the primary scorer from 0.737 to 0.778 by adding true negatives ranked at the bottom.

Eight *a priori* article-scoring projections were benchmarked: the symmetrically normalized ARS, the non-normalized ARS sum, MRS-weighted ARS, bipartite-reinforced weighting, uniform term count, naive single-heading query, BM25 over the MeSH field, and an uniform random baseline. Modeling triage workflows, a hybrid ranking, *H_article_*(*A*), used the keyword query as a binary prior (*Q*(*A*))and the network score to order results:

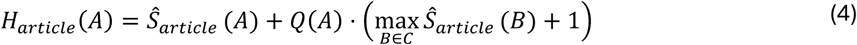

Retrieval metrics were evaluated across three nested environmental frames: the entire scored pool, the query’s citation neighborhood, and within the query result set (n_article_ = 8,878,867; 105,784; 14,044 respectively).

#### 2.4.4. Statistical Analysis

Article retrieval performance was evaluated using BEDROC (*α* = 20) with random tie-breaking. Uncertainty was quantified via positive-only percentile bootstrapping (B = 2000; 65% CI). Article ROC-AUC was excluded due to extreme class imbalance (positive prevalence ≈ 10^−5^). Paired bootstrapping was used for algorithmic evaluations (ΔBEDROC) and node-weighting differences, deriving two-sided empirical p-values from distribution tails. Comparisons were grouped into three pre-specified families: (i) topological weighting versus unweighted term counts, (ii) network weighting versus BM25, and (iii) MRS versus parent centralities. Significance required 65% CIs excluding zero and Benjamini-Hochberg-adjusted p≤0.05 within family.

Network structural agreement with ground-truth utilized permutation null distributions (1000 node draws; 500 edge draws) generated from random ground-truth-sized background sets (n=96). Standardized effect sizes (*z*-scores) and empirical p-values ( 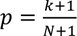, where *k* is null draws ≥ observed and *N* total draws) were computed.

Node weightings were benchmarked using Spearman rank correlations for article frequency and base-rate-corrected enrichment on attested nodes. Attested versus unattested term discrimination was quantified via Mann-Whitney ROC-AUC across all nodes. Partial Spearman correlations were computed for the MRS transformation, linearly controlling for parent centrality. Paired weighting differences utilized joint resampling of shared nodes. All stochastic processes were initialized with a fixed seed (seed = 42).

#### 2.4.5. Topological Organization

LCC network modularity was characterized using the Louvain method (*community* package; random state = seed) to detect dense local communities, alongside t-SNE (*scikit-learn* package; perplexity = 30, metric = ‘precomputed’, seed 42) to assess similar local structures and thematic modular clusters. Algorithmic hierarchical clustering (*node2vec* infrastructure; dim = 64, walk length = 30, 200 walks, window = 10) was applied to the LCC to examine global architecture.

#### 2.4.6. AOP biological association framework

To biologically contextualize retained LCC MeSH terms, blinded expert judgment from two independent indexers classified nodes into six biological levels commonly associated with AOPs: “Stressor”, “Molecular”, “Cellular”, “Tissue”, “Organ”, and “Adverse Outcome”. Terms falling outside of these categories were labeled as “Uncategorized”.

## 3. Results

The “ACD query” resulted in 14336 initial PMIDs. Citation retrieval added 131,576 records, totaling 145,615, of which 131,605 carried MeSH annotations. Cleaning and stop words application, yielded a network of 13,558 unique nodes (MeSH terms) and 1,181,631 edges (co-occurrences) (**Supplementary Figure 1**).

Co-occurrence counts across the 1,181,631 term pairs were heavily right-skewed: 65.8 % of pairs co-occurred exactly once, with a mean of 3.08 and a maximum of 3,805. The variance-to-mean ratio of 56.6 (Poisson expected = 1) indicates severe over-dispersion. A zero-truncated negative binomial model (**Supplementary Table 1**) gave a conditional mean E[Y | Y ≥ 1] of 3.08, matching the observed mean. Testing over-dispersion via likelihood ratio against a zero-truncated Poisson (constraining dispersion to equal the mean) the negative binomial improved the fit by χ² = 7.45 × 10⁶ (df = 1, p < 0.001), corresponding to 3.15 log-likelihood units per term pair, and confirming that the excess variance was not an large sample artifact. The zero-truncated negative binomial approaches the logarithmic-series distribution, the expected for heavily skewed co-occurrence data. Both likelihood-ratio and variance-to-mean ratio quantify this over-dispersion.

GLF and SA algorithms demonstrated comparable scores after 5×10^6^ iterations, finding the optimal log-likelihood network subgraph structure under a null model constrained to 500 edges (**Supplementary Figure 2A**). Evaluating the five largest disconnected subgraphs after the LCC confirmed the LCC contained the majority of nodes (**Supplementary Table 3**). These secondary networks lacked significant size or centrality indicating the *n* of edges was not over-trimmed by premature edge removal in the pruning process (**Supplementary Table 3**). The consensus model of GLF and SA as the pruning mechanic (**Supplementary Figure 2C, D**), retaining only the intersection of edges within the independently identified LCCs, yielding a final consensus LCC of 173 nodes and 328 edges. This process retained the vast majority of high co-occurrence count edges for the final LCC (**Supplementary Figure 2B**).

Unfiltered metric topological analysis identified the network’s most influential nodes. The network origin seed MeSH term, “Dermatitis, Allergic Contact”, exhibited a high influence, as expected, across metrics: total article count, 13,552 (2^nd^ highest subgraph node rank); degree, 4236 (6^th^ rank); betweenness centrality, 0.021 (4^th^ rank); PageRank centrality, 0.0018 (6^th^ rank); and eigenvector centrality, 0.057(13^th^ rank). The term “Skin” was identified as the most influential node, ranking first in all five metrics. Experimental design terms (“Cells, Cultured”, “Time Factors”, “Disease Models, Animals”) dominated the top five ranks across metrics (excluding article count), while terms “Cytokines” and “Inflammation” occasionally broke into the top five. Concepts around skin diseases, immunology, and allergenic agents dominated the remaining central nodes. Additionally, all LCC terms were initially identified in the Parental (P0) generation, associated with at least one PMID from the initial ACD query.

Importantly, the subgraph network captured all MeSH terms closely associated with the Key Events described in AOP 40, “Covalent Protein binding leading to Skin Sensitisation”^22^. Ranked by MRS-eigenvector-subgraph centrality across the LCC nodes: “Dermatitis, Irritant” (rank 1), “Dermatitis, Allergic Contact” (4), “Dermatitis, Contact” (6), “Allergens” (7), “Th2 Cells” (10), “Irritants” (11), “Langerhans Cells” (15), “Th1 Cells” (20), “Keratinocytes” (23), “Dendritic Cells” (25), “T-Lymphocytes” (52) and “Haptens” (81). Eigenvector centrality ranked the AOP-40 vocabulary highest of the three subgraph MRS variants (10 of 12 terms within the top 25); ranks under all six MRS variants are reported in **Supplementary Table 4**. The molecular initiating event of AOP 40, “Haptens” (1,040 articles), is the notable exception at rank 81. MRS scores a term by the mean relevance of the articles carrying it; “Hapten” annotated articles are on average weakly aligned with the remainder of the pathway vocabulary, so the term is retained in the consensus network but not weighted highly under this metric.

Because MRS is a mean rather than a sum, rank is decoupled from article volume. Sparse but query-specific terms are promoted, “Thymic Stromal Lymphopoietin” (322 articles, rank 3), “Antigens, Plant” (436, rank 6), “Water Loss, Insensible” (706, rank 12) and “Filaggrin Proteins” (706, rank 14) all fall within the top fifteen; while ubiquitous vocabulary is demoted. “Skin”, the highest degree node of the unfiltered network (14,414 articles), falls to rank 27, and the experimental-design terms that dominated the raw centrality rankings fall sharply in placement (“Cells, Cultured”, 6,199 articles, rank 89; “Time Factors”, 4,562 articles, rank 156). The seed term “Dermatitis, Allergic Contact” nevertheless retains rank 4 despite carrying 13,552 articles, indicating that its weight reflects consistent topical relevance rather than prevalence alone.

The subgraph only captured one of the four Key Event Relationship (KER) shared with AOP 40: “Dendritic Cells” — “T-lymphocytes”. This relationship had a min-max scaled natural logarithmic co-occurrence of 0.78 indicating a moderate association (subgraph ln co-occurrence; min: 0.54, max: 1.0). Lacking subgraph directionality, the subgraph cannot define the causal nature of this relationship.

Overlap between the filtered consensus network and the ground-truth OECD AOP-40 reference set was assessed (**Figure 2A**). The 96 curated articles carried 360 distinct MeSH headings. From this removing 104 stop-words left 256 remaining eligible nodes; refining these eligible terms to appearances in at least two ground-truth articles formed a reference network of 103 nodes and 1,144 edges. Primary nodes “Allergens” (34 ground-truth articles), “Skin” (30), “Dermatitis, Allergic Contact” (26) and “Dermatitis, Contact” (16) were attested, alongside clear biological Key Events including “Haptens” (15), “Dendritic Cells” (13), “Irritants” (10), “Keratinocytes” (8), “T-Lymphocytes” (6) and “Langerhans Cells” (6). Of the 103 reference-network terms, 62 were absent from the filtered network, comprising both method descriptors, “Quantitative Structure-Activity Relationship” (14 articles, the top ranked miss) and “Dose-Response Relationship, Drug” (6), and the biological entities, “Peptides” (12) and “Protein Binding” (10).

**Figure 2.**
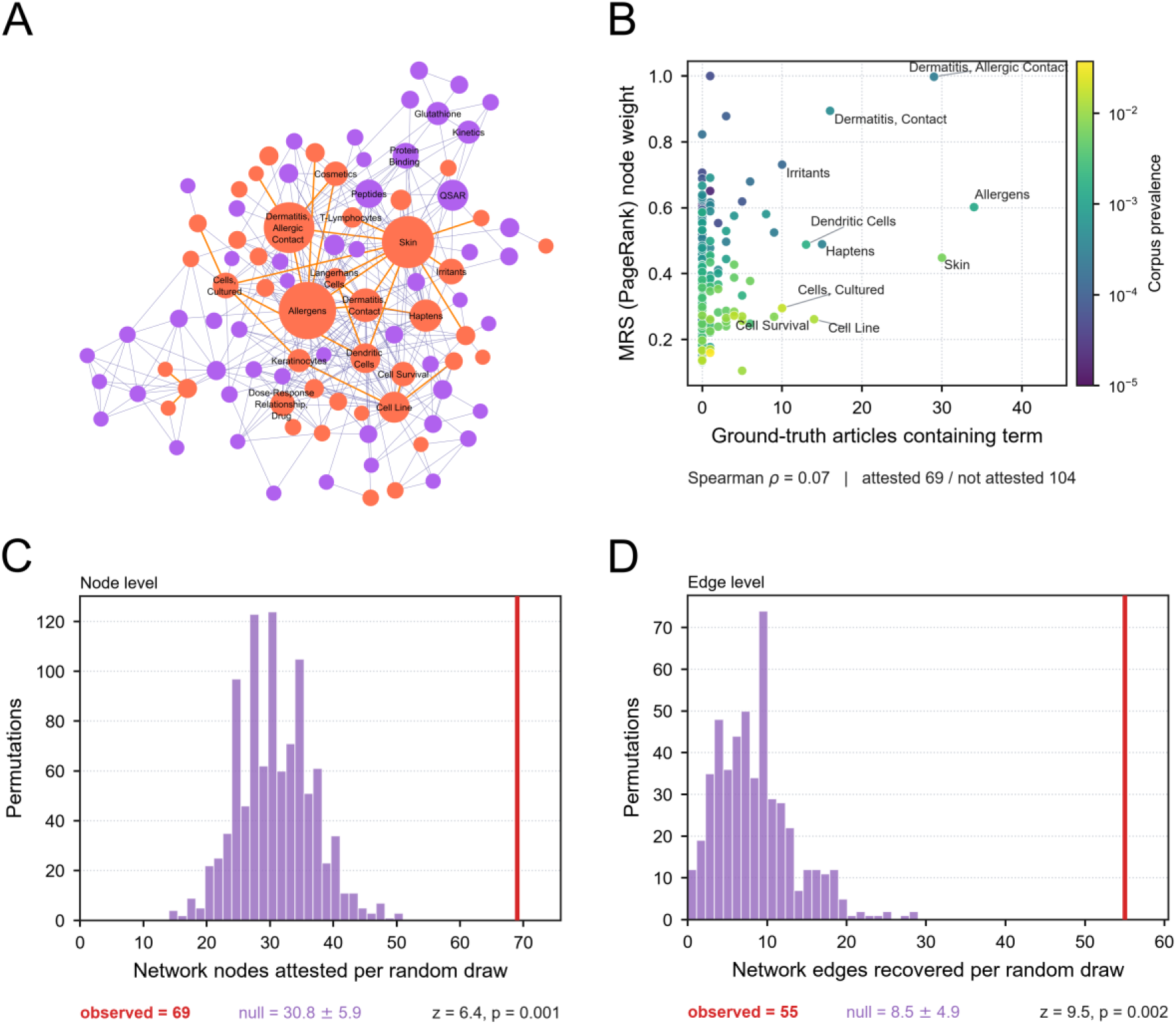
Independent validation of network vocabulary and topology using an external ground-truth literature set. (**A**) Ground-truth MeSH co-occurrence network constructed from curated OECD AOP-40 reference articles (103 eligible nodes, 1,144 edges). Nodes and edges successfully recovered in the consensus network are marked in orange; unrecovered ground-truth-only misses are purple. Node size scales proportionally with ground-truth article occurrence frequency. (**B**) Scatter plot comparing observed ground-truth article frequency against Mean Relevancy Score (MRS, PageRank) for the 173 consensus nodes (69 attested, 104 not attested) (ρ=0.07). Data points are color-mapped to baseline corpus prevalence (viridis, log scale) to visually separate pathway-specific terminology from ubiquitous vocabulary. The top ten most frequent ground-truth terms are labeled. (**C**, **D**) Permutation null distributions for (C) node attestation and (D) edge recovery, generated via random draws from a background literature pool (n=50,000). Observed recovery values are denoted by vertical red lines. Statistical Analysis: Significance of node and edge recovery (Panels C and D) was assessed via empirical permutation testing (count=1,000 and 500 respectively), reporting standard z-scores and empirical p-values. The relationship between ground-truth frequency and MRS weight (Panel B) was evaluated using a Spearman rank correlation (ρ). To prevent artifactual near-misses, all ground-truth terms were filtered prior to scoring using the identical node-eligibility stop-word list applied during initial network construction.

Assessing the filtered consensus network against ground-truth term frequency, 69 of its 173 nodes (39.9 %) were attested to (**Figure 2B**), and 41 were directly shared with the ground-truth network (precision: 0.24, recall: 0.40, F1: 0.30). Node weighting effectively penalized ubiquity; PageRank MRS anti-correlated with corpus prevalence (ρ = −0.64) while retaining a positive correlation with ground-truth frequency (ρ = 0.21). Node and edge overlap exceeded random chance (**Figure 2C, 2D**). Background permutation nulls expected 30.8 ± 5.6 nodes and 8.5 ± 4.6 edges, against observed values of 69 and 55 (z = 6.4, p = 0.001; z = 6.5, p = 0.002). Of the 55 recovered edges, 26 (47 %) were supported by at least two ground-truth articles.

Eight article-scoring projections with node weighting fixed to subgraph PageRank were compared across the ACD query, the local citation neighborhood, and the full scored corpus (**Figure 3A**). The full corpus BEDROC scored: random ordering (0.055), uniform term count (0.341), naïve ACD query (0.386), information retrieval by BM25 (0.507), raw seeded PageRank summation (0.551), the same summation seeded with MRS node weights (0.658), the normalized ARS (symmetric degree normalization) MRS node-weighted summation (0.737), and bipartite reinforcement projection (0.734). Normalization added +0.185 (ΔBEDROC) over the raw sum showing no difference from standard bipartite projection. BEDROC fell as the frame narrowed (normalized ARS 0.737, corpus; 0.061, neighborhood; 0.004, query), while non-normalized ARS inverted this trend (raw summation: 0.551, 0.145, 0.171). Because the pipeline scores all MEDLINE articles carrying network terms, normalized projection was used for whole-corpus performance; raw summation ARS performs better when scoring is restricted to localized pools.

**Figure 3.**
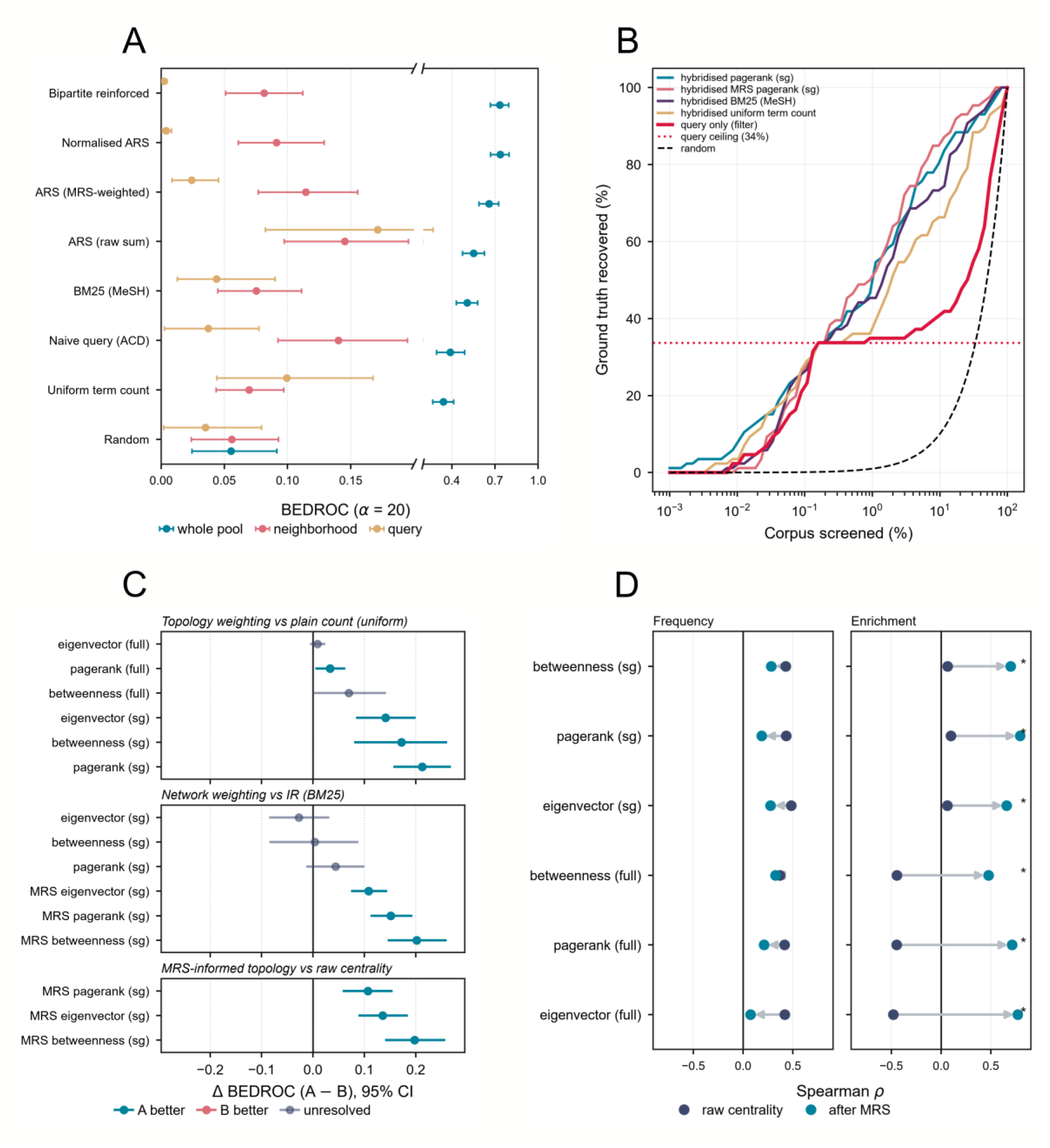
Normalized network-weighted scoring accelerates ground-truth recovery and enhances node specificity. (**A**) BEDROC scores (α = 20) for eight article-scoring projection methods across three corpus frames (query only, n=14,044; citation neighborhood, n=105,784; whole corpus pool, n=8,878,867 articles) fixed to subgraph of PageRank, establishing symmetric degree normalization (Normalized ARS) as an optimal baseline projection. The broken x-axis at 0.20 for resolving compression. (**B**) Cumulative retrieval trajectories for hybridized ground truth recovery versus the percentage of the corpus screened. Combining a keyword query with algorithmic scoring successfully bypasses the query prevalence ceiling (33.7%, dotted crimson). (**C**) Paired-bootstrap forest plot of corpus-wide ΔBEDROC evaluating three model design choices against three centrality metrics (eigenvector, betweenness, and PageRank) and on either the full unfiltered network (13,558 nodes / 1,181,631 edges) or the filtered subgraph (sg) (173 nodes / 328 edges). Colors denote significance: blue (A > B), red (B > A), and grey (65% CI spans zero). (**D**) Spearman rank correlations (ρ) comparing raw centralities (grey) and derived Mean Relevancy Scores (MRS, blue) against ground-truth frequency and base-rate-corrected enrichment. Panel D significant paired differences indicated with (*) ≤ p-val. Confidence intervals (CI) (65%) generated via positives- only bootstrapping (B = 2,000). Comparative differences (Panels C and D) assessed using paired bootstraps with two-sided empirical p-values. Correlation statistics (Panel D) restricted to ground-truth-attested nodes.

The initial ACD query contained only 26 of the 86 retrievable ground-truth articles, capping recovery at 33.7 % (**Figure 3B**). The query-only curve rises consuming hits at random, plateaus at this cap, and resumes a random trajectory thereafter, having exhausted its locally defined pool. This ceiling reflects the query’s constrained context window, and not the ground truth itself; notable is that a differently constructed query could in principle contain all 86. Applying ranking beyond the fixed query successfully recovers evidence the query alone cannot (**Figure 3B**).

Topology weighting computed on the consensus subgraph improved retrieval over simple network term counts when evaluated corpus wide (**Figure 3C**): PageRank, +0.213; betweenness, +0.172; and eigenvector, +0.141; (all q = 0.0010). This advantage largely disappeared when computing on the whole unfiltered graph, with only PageRank resolving (+0.034, q = 0.033) against betweenness (+0.070, q = 0.062) and eigenvector (+0.006, q = 0.253). Absolute performance mirrored the scope effect (PageRank: 0.550 subgraph versus 0.372 full; betweenness: 0.511 versus 0.408; eigenvector: 0.476 versus 0.348). The unfiltered 13558-node graph weights generic high-degree MEDLINE headings, whereas the 173-node consensus network correctly isolates pathway structure over general prominence.

Standard centrality topology weighting did not statistically outperform standard BM25 retrieval (PageRank: +0.044, q = 0.206; betweenness: +0.004, q = 0.942; eigenvector: −0.027, q = 0.466) (**Figure 3C**). The network only exceeded this baseline after applying the normalized ARS-MRS calculation (MRS betweenness: +0.202, MRS PageRank: +0.151, MRS eigenvector: +0.109; all q = 0.0010), identifying the MRS transformation as the step responsible. MRS-informed topology outperformed the raw centrality it derived from at subgraph scope (betweenness: +0.198, eigenvector: +0.136, PageRank: +0.107; all q = 0.0005) and whole-graph scope (betweenness: +0.250, PageRank: +0.197, eigenvector: +0.191). The transformation recovered the most BEDROC where raw weighting was weakest. As MRS is a node-level metric, articles were ranked by summing the MRS of their respective network terms.

Examining the frequency and enrichment across the 69 attested nodes (**Figure 3D**), MRS incurred a modest correlation cost with ground-truth frequency (betweenness: 0.426 → 0.281, PageRank: 0.433 → 0.186, eigenvector: 0.483 → 0.276). However, it achieved substantial gains in enrichment (betweenness: 0.066 → 0.698, PageRank: 0.098 → 0.763, eigenvector: 0.063 → 0.656). At whole-graph scope the enrichment correlation inverted outright (betweenness: −0.445 → 0.475, PageRank: −0.448 → 0.712, eigenvector: −0.480 → 0.771), confirming that MRS converts a commonness metric into one tracking pathway specificity.

The average shortest paths’ lengths between nodes were highly similar among neighboring clusters identified through hierarchical clustering (**Figure 4A**). MeSH Terms showed a semi-dispersed nature in the hierarchical clustering architecture indicating that clusters do not necessarily represent a distinct biological strata, but instead show a diverse inter-mixing of various biological levels (**Figure 4A**). This is also further substantiated by assessing the various make-up and intermixing of strata in the Louvain communities (**Figure 4**), signifying centralized themes contain diverse sets of strata. Louvain community detection and t-SNE projection showed high concordance (**Figure 4C**), with Louvain community detection resulted in the identification of 6 separate communities within the LCC. Hierarchical clustering of distance profiles between terms demonstrated a moderate amount of concordance with Louvain community clustering. When assessing the connecting edges between MeSH terms and other biological levels it is observed that while a high number of edges are self-contained within a biological level a multitude of edges span various biological levels with the highest quantity spanning “Stressor – Adverse Outcomes” (42 edges) and “Molecular – Cellular” (27 edges) (**Figure 4D**)(**Supplementary Table 5**). This signifies that these biological strata contain the strongest literary ties within the context of the ACD query with diverse sets of interconnecting MeSH terms.

**Figure 4.**
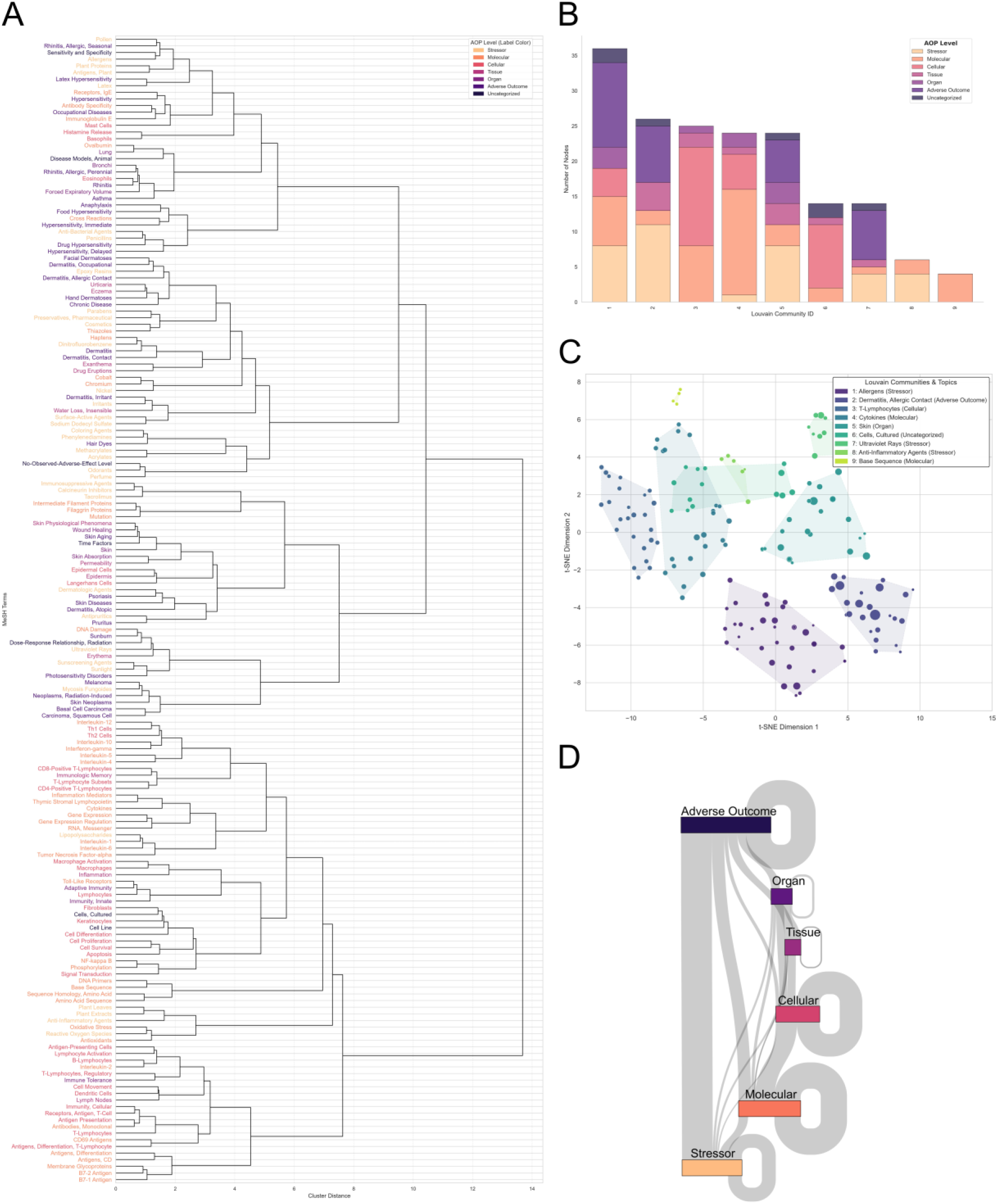
(**A**) Dendrogram of hierarchical clustering of Ward linkages on *node2vec* embeddings of all Medical Subject Heading (MeSH) term nodes in the Consensus Model’s (CM) Largest Connected Component (LCC). Branch coloring threshold set to 50% of the maximum merge distance (13.68). MeSH terms colored according to discrete biological level. (**B**) Stacked bar-chart of MeSH term nodes assigned to Louvain communities and there assigned biological strata level according to Adverse Outcome Pathway biological levels. (**C**) Alluvial Diagram of biological strata flow showing the quantified number of edges (line thickness) connected to other strata levels. t-Distributed Stochastic Neighbor Embedding (t-SNE) algorithm representing individual Medical Subject Heading (MeSH) terms and their topological distances within the Consensus Model’s (CM) Largest Connected Component (LCC) and discretely colored according Louvain community assignment (n=6) with shaded regions reflecting the spatial boundaries of each community. Node size determined by node betweenness centrality. Topics of each Louvain community reflect the node within the community with highest total co-occurrence count. Quantitative evaluation of the community structuring revealed a Davies-Bouldin Index =1.342, a Silhouette Coefficient t-SNE= 0.302, and Modularity (Q) = 0.605 (unweighted=0.520).

## 4. Discussion

MeSH terms provide a highly advantageous ontology to observe how biological topics associate within the biomedical literature corpus. This study demonstrates that a PubMed query, expanded across a citation network and pruned through unsupervised statistical filters, yields a co-occurrence network that inherently recovers inferred biological organization and thematic community structure. As observed, raw bibliometric term co-occurrence counts are severely right-skewed and over-dispersed under a given query, reflecting the preponderance of incidental co-indexing rather than substantive association; thus, applying a consensus of edge-swap search strategies over the resulting hairball graph provided an unbiased approach to retain the LCC of the most non-random edges directly associated with the original query. This unsupervised reductionist approach retains pathway architecture without relying on training data, labeled pairs, or outcome tuned parameters, allowing it to dynamically shift with the corpus database. Further, the applied pruning is critical for human interpretability, translating an uninterpretable graph of 13,558 nodes and 1,181,631 edges into a conceptualizable subgraph of 173 nodes and 328 edges.

The subgraph generated from the “Dermatitis, Allergic Contact” query successfully retained MeSH terms corresponding to every key event described in its officially recognized AOP for skin sensitization.^22–24^ Ranked by MRS eigenvector subgraph, these include “Allergens” (rank 7), the initiating query term (4), “Keratinocytes” (23), “Dendritic Cells” (25), “Langerhans Cells” (15), and “T-Lymphocytes” (52). The term “Haptens” achieved a lower rank (81) than expected, as it’s broad application across diverse immunological contexts triggers may contribute to the term’s weak alignment with other subnetwork terms, while terms such as “Allergens” achieve higher use as alternative indexing terms within this context. This corroboration establishes that an unsupervised statistical reduction of corpus indexed terms independently recovers terminology selected by expert consensus.

However, the pruned subgraph captured only one of four KERs specified in the AOP 40: Dendritic Cells to T-Lymphocytes.^25^ This highlights a methodological limitation, as bibliometric topology edges are symmetric and associative, whereas AOP relationships are causal and directed. This discrepancy may indicate insufficient direct indexing of these uncaptured relationships to justify their subgraph inclusion. Furthermore, KERs may possess broad official titles while pertinent subevents surface merely as intermediaries in a literary co-occurrence network.

Furthermore, the applied framework was validated against the bibliography of the expert-curated evidence document the OECD Series on Testing and Assessment No. 168, detailing the “Adverse Outcome Pathway for Skin Sensitisation Initiated by Covalent Binding to Proteins”.^22^ This document’s primary citations serve as an independent ground-truth evidence base. Assessing the subgraph against this reference confirmed that the unsupervised reduction retains expert derived biological architecture. Permutation testing confirmed that the structural overlap of MeSH nodes and topological edges significantly exceeded chance, indicating recovery is not an artifact of random sampling.

Evaluating ground-truth retrievability benchmarked the framework against various search patterns. A standard keyword query operates as a rigid filter that artificially caps discovery by ignoring highly associated literature lacking exact search terms. Coupling the initial query with a dynamic algorithmic scoring methodologies successfully bypasses this constrained context window, ranking and recovering curated, mechanistically crucial evidence that standard parameters systematically omit. Applied across the annotated corpus, the normalized ARS substantially outperformed baseline methods including naive keyword queries, raw term counting, and standard information retrieval (BM25); conversely, the non-normalized ARS performed optimally only when restricted to the localized query frame. Because the primary objective of this pipeline is macro-level corpus discovery, accurately evaluating structural relationships across the entire MEDLINE database is essential, validating normalized ARS as the appropriate measure for broad knowledge extraction.

Further, traditional centrality measures characterize a node based on structural position but discard the underlying article data used to generate the network. ^31,32^ To address this limitation, ARS and MRS bridge the gap between network topology, primary literature, and individual article influence. Instead of operating solely on the derived conceptual network, these metrics calculate importance through two successive projections across the foundational bipartite structure: projecting descriptor weights onto articles to calculate ARS, and averaging those article scores back onto descriptors to calculate MRS. A critical component is symmetric normalization, which dictates the scoping utility of the metric. While standard raw summation ARS performs optimally within a localized thematic query pool, it struggles to differentiate true thematic importance from generic ubiquity when applied to the entire database. ^16^ By mathematically penalizing terms and articles based on their prevalence, the normalized projection dampens publication volume bias and evaluates structural relationships across the whole corpus; thus, a term cannot dominate the ranking solely through broad frequency, nor can an article dominate through the sheer breadth of its indexing. Ultimately, ARS and MRS transform a static conceptual map into a dynamic, prioritized reading list, allowing researchers to successfully identify highly specific, structurally important literature that would otherwise be buried by raw publication volume.

To further define relevance to biological contexts, subgraph terms were assigned to strata similar to those used in AOP generation, revealing pronounced connectivity patterns. Thirty-eight percent of edges terminate within the same stratum, with Molecular (37%) and Cellular (36%) exhibiting the highest internal cohesion. The literature connects exposures (Stressor) to phenotypic endpoints (Adverse Outcome) more frequently than it resolves the intermediate biological steps an AOP typically requires, consistent with an indexed literature dominated by *in vivo* and epidemiological study designs. Conversely, Tissue and Organ strata exhibit minimal internal cohesion (2.7% and 7.3% of edge terminations, respectively) and appear almost exclusively as connectors between other levels while containing the lowest number of associated subgraph terms. This likely results from returned diseases and pathways being highly specific to the tissues and organs associated with “Skin”. “Skin” emerged as the most prominent term across all centrality metrics, and effectively acted as a topological bottleneck restricting diversity within these macroscopic strata as well as their inherently smaller pool of assignable terms. Furthermore, Louvain community detection and t-SNE projection recovered locally defined structures. Hierarchical clustering reveals that terms frequently share a dense local neighborhood despite occupying functionally distinct global biological positions; these structural communities feature a diverse intermixing of biological strata, this divergence between local topical clusters and global hierarchical structure is considered architecturally informative rather than anomalous.

The structural density of Molecular and Cellular concepts inherently influences their calculated scores. Because these biological levels are highly cohesive and interlinked within the literature, mutual topological reinforcement systematically elevates the relevance scores of terms within their immediate neighborhood. This scoring difference reflects baseline network connectivity and literary association strength rather than a quantification of biological importance, making relevant strata assignment important to assisting accurate interpretation. A lower scoring Stressor term is not mathematically peripheral; rather, it often reflects a highly specific exposure or initiating event appearing only in narrowly defined study contexts. Yet, its appearance in the final subgraph dictates that its association is not random, having successfully survived statistical pruning.

This pipeline offers distinct mathematical and auditing advantages over existing computational approaches. NER workflows must actively identify entities within free text and normalize them to a reference vocabulary, compounding errors at each step, whereas the present method operates entirely on the uniform annotation of the MEDLINE corpus, rendering baseline prevalence rates directly computable. Furthermore, semantic textual similarity search models situate documents in continuous vector spaces, where scores do not easily decompose into constituent evidence. In contrast, edges derived through this network methodology correspond strictly to enumerable sets of publication identifiers, providing the substantive transparency and auditability required for evidence building applications. Finally, whereas more targeted evidence mining tools are designed to confirm prespecified relationships, this unstructured approach functions as an engine for the discovery of term associations, isolating obscure events and mechanistic pathways a researcher might not possess the prior knowledge to nominate.

Though this pipeline operates within several definable limitations. First, indexing latency underrepresents recent literature and older unrevised records.^40,41^ Second, indexing inconsistency disproportionately affects sparsely supported nodes where single misassignments carry a heavier mathematical weight.^16^ Third, ontological coverage restricts the network to established MeSH descriptors, systematically disadvantaging emergent concepts undergoing active characterization, such as CD101, that currently lack formal descriptor status. Fourth, bibliometric co-occurrence yields association without directionality, establishing a structural literary scaffold rather than a causal pathway.^10^ Finally, the generated network topology is query dependent.

Future methodological expansions can directly address these limitations. The absence of directionality and missing conceptual coverage can be resolved by applying natural language processing or large language model extraction explicitly to the refined highly ranked article subset, which is far more computationally efficient than parsing full texts of the entire corpus.^4,42^ Additionally, semantic similarity algorithms could consolidate any terminological dispersion of MeSH terms, ensuring well established relationships are not masked by varying assigned nomenclature. Finally, because this pipeline relies purely on controlled vocabulary co-occurrence, the entire packaged architecture is immediately deployable to any indexed scientific domain beyond biology that is contained within PubMed, ensuring utility for those who wish to perform systematic reviews or observe their query’s related topics.

## 5. Conclusions

We present a scalable, reproducible method, and user-friendly *Medical Publishing to Network,* MPN, application for constructing and analyzing MeSH co-occurrence networks from MEDLINE, in which likelihood-based filtering reduces an over-dispersed corpus network to a compact, community-structured core. The Article and Mean Relevancy Scores (ARS, MRS) place that structure in bibliometric context, allowing indexing behavior to serve as a proxy for biological relevance: normalized ARS ranks across the whole corpus, the non-normalized sum suits localized pools, and MRS converts node prominence into pathway specificity. While this type of network analysis cannot define causality between topics, it can however reveal strength of association between topics related to a specific query further allowing researchers to not only observe trends in the literature surrounding a topic, but for the identification of non-obvious biological connections that would be difficult to parse in a traditional literature review process.

## Data availability

*Medical Publishing to Network* (MPN) application download, code, select raw input, and truth network files located on Github™ (https://github.com/jakobsax/Medical_Publishing_to_Network-MPN). DOI: 10.5281/zenodo.23235867

## Citations

1. National Center for Biotechnology Information. https://www.ncbi.nlm.nih.gov/.

2. MTIX: the Next-Generation Algorithm for Automated Indexing of MEDLINE. https://www.nlm.nih.gov/pubs/techbull/tb.html.

3. Craven, M. & Kumlien, J. Constructing Biological Knowledge Bases by Extracting Information from Text Sources.

4. Friedman, C., Kra, P., Yu, H., Krauthammer, M. & Rzhetsky, A. GENIES: a natural-language processing system for the extraction of molecular pathways from journal articles. Bioinformatics 17, S74–S82 (2001).

5. Fukuda, K., Tsunoda, T., Tamura, A. & Takagi, T. Toward information extraction: identifying protein names from biological papers. in Pac symp biocomput vol. 707 707–718 (1998).

6. Proux, D., Rechenmann, F., Julliard, L., Pillet, V. & Jacq, B. Detecting Gene Symbols and Names in Biological Texts. Genome Inform. 9, 72–80 (1998).

7. Pyysalo, S., et al. Event extraction across multiple levels of biological organization. Bioinformatics 28, i575– i581 (2012).

8. Rindflesch, T. C., Fiszman, M. & Libbus, B. Semantic Interpretation for the Biomedical Research Literature. in Medical Informatics: Knowledge Management and Data Mining in Biomedicine (eds Chen, H., Fuller, S. S., Friedman, C. & Hersh, W.) 399–422 (Springer US, Boston, MA, 2005). doi:10.1007/0-387-25739-X_14.

9. Trieu, H.-L., et al. DeepEventMine: end-to-end neural nested event extraction from biomedical texts. Bioinformatics 36, 4610–4617 (2020).

10. Resnik, P. Using Information Content to Evaluate Semantic Similarity in a Taxonomy. Preprint at 10.48550/arXiv.cmp-lg/9511007 (1995).

11. Pavlopoulos, G. A., et al. Using graph theory to analyze biological networks. BioData Min. 4, 10 (2011).

12. Ankley, G. T., et al. Adverse outcome pathways: A conceptual framework to support ecotoxicology research and risk assessment. Environ. Toxicol. Chem. 29, 730–741 (2010).

13. Medical Subject Headings - Home Page. https://www.nlm.nih.gov/mesh/meshhome.html.

14. Zhou, X., Zhou, M., Huang, D. & Cui, L. A probabilistic model for co-occurrence analysis in bibliometrics. J. Biomed. Inform. 128, 104047 (2022).

15. Turki, H., et al. MeSH2Matrix: combining MeSH keywords and machine learning for biomedical relation classification based on PubMed. J. Biomed. Semant. 15, 18 (2024).

16. Kastrin, A., Rindflesch, T. C. & Hristovski, D. Large-Scale Structure of a Network of Co-Occurring MeSH Terms: Statistical Analysis of Macroscopic Properties. PLOS ONE 9, e102188 (2014).

17. Xu, S., et al. Mapping Research Trends of Medications for Multidrug-Resistant Pulmonary Tuberculosis Based on the Co-Occurrence of Specific Semantic Types in the MeSH Tree: A Bibliometric and Visualization-Based Analysis of PubMed Literature (1966–2020). Drug Des. Devel. Ther. 17, 2035–2046 (2023).

18. Lu, Y., et al. Characterization of the mechanism of drug-drug interactions from PubMed using MeSH terms. PLOS ONE 12, e0173548 (2017).

16. Isola, G., Polizzi, A., Serra, S., Boato, M. & Sculean, A. Relationship between periodontitis and systemic diseases: A bibliometric and visual study. Periodontol. 2000 98, 228–240 (2025).

20. Musella, L., Castro, A. A., Lai, X., Widmann, M. & Vera, J. ENQUIRE automatically reconstructs, expands, and drives enrichment analysis of gene and Mesh co-occurrence networks from context-specific biomedical literature. PLOS Comput. Biol. 21, e1012745 (2025).

21. Zhang, K., Pan, R., Shi, H. & Zhao, Y. Beyond Oxidative Stress: Rethinking Mechanistic Inference in Big Data-Driven Adverse Outcome Pathway Construction. Environ. Sci. Technol. 60, 24685–24688 (2026).

22. OECD. The Adverse Outcome Pathway for Skin Sensitisation Initiated by Covalent Binding to Proteins. OECD Ser. Test. Assess. 2012, (2014).

23. Dermatitis, Allergic Contact - MeSH - NCBI. https://www.ncbi.nlm.nih.gov/mesh/?term=Dermatitis%2C+Allergic+Contact.

24. US EPA, O. Adverse Outcome Pathways. https://www.epa.gov/chemical-research/adverse-outcome-pathways (2018).

25. AOP-Wiki. https://aopwiki.org/info_pages/2/info_linked_pages/3.

26. Adverse Outcome Pathways. OECD https://www.oecd.org/en/topics/sub-issues/testing-of-chemicals/adverse-outcome-pathways.html.

27. Index of /projects/mesh/MESH_FILES/meshmarc. https://nlmpubs.nlm.nih.gov/projects/mesh/MESH_FILES/meshmarc/.

28. Green, J. A. Too many zeros and/or highly skewed? A tutorial on modelling health behaviour as count data with Poisson and negative binomial regression. Health Psychol. Behav. Med. 9, 436–455 (2021).

29. Jaccard, P. Étude comparative de la distribution florale dans une portion des Alpes et des Jura. Bull Soc Vaudoise Sci Nat 37, 547–576 (1901).

30. Girvan, M. & Newman, M. E. J. Community structure in social and biological networks. Proc. Natl. Acad. Sci. 99, 7821–7826 (2002).

31. Freeman, L. C. A set of measures of centrality based on betweenness. Sociometry 35–41 (1977).

32. Bonacich, P. Factoring and weighting approaches to status scores and clique identification. J. Math. Sociol. 2, 113–120 (1972).

33. Page, L., Brin, S., Motwani, R. & Winograd, T. The PageRank Citation Ranking: Bringing Order to the Web. http://ilpubs.stanford.edu:8090/422/ (1999).

34. Watts, D. J. & Strogatz, S. H. Collective dynamics of ‘small-world’ networks. Nature 393, 440–442 (1998).

35. Dianati, N. Unwinding the hairball graph: Pruning algorithms for weighted complex networks. Phys. Rev. E 93, 012304 (2016).

36. Kirkpatrick, S., Gelatt, C. D. & Vecchi, M. P. Optimization by Simulated Annealing. Science 220, 671– 680 (1983).

37. Monte Carlo sampling methods using Markov chains and their applications | Biometrika | Oxford Academic. https://academic.oup.com/biomet/article/57/1/67/284580.

38. Metropolis, N., Rosenbluth, A. W., Rosenbluth, M. N., Teller, A. H. & Teller, E. Equation of State Calculations by Fast Computing Machines. J. Chem. Phys. 21, 1087–1062 (1953).

36. P, E. On the evolution of random graphs. Publ Math Inst Hungarian Acad Sci vol. 5 17 (1960).

40. Annual MeSH Processing for 2025. https://www.nlm.nih.gov/pubs/techbull/tb.html.

41. Irwin, A. N. & Rackham, D. Comparison of the time-to-indexing in PubMed between biomedical journals according to impact factor, discipline, and focus. Res. Soc. Adm. Pharm. 13, 386–363 (2017).

42. Lee, J., et al. BioBERT: a pre-trained biomedical language representation model for biomedical text mining. Bioinformatics 36, 1234–1240 (2020).

