## Supplementary Materials for "Bridging Bibliometrics and Biology: A Framework for the Relevance Scoring and Biological Stratification of Scalable MeSH Co-occurrence Networks"

**Supplementary Note 1 - Consensus Modeling**

The null model assumes that the probability of a co-occurrence between any two nodes (MeSH terms) is proportional to the product of their strengths. Let $G = (V, E)$ represent a given network. The strength ($s_i$) of node *i* is defined as the sum of weights, w, of its edges with adjacent node. With $W_{total}$ representing the collective weight of the full network:

$$s_i = \sum_j w_{ij} \qquad\qquad W_{total} = \sum_{(i,j)\in E} w_{ij} \qquad \text{(S1),(S2)}$$

Where $w_{ij}$ is the observed co-occurrence count for the edge between nodes *i* and *j*. The total strength of the graph, $W_{total}$, is the sum of all edge weights in the set of all unique edges in the network, *E*. Node strengths and $W_{total}$ were computed once on the full unfiltered network and held fixed throughout the search, such that all candidate subgraphs are evaluated against a single global null. Since $\Sigma i\ s_i = 2W_{total}$, the factor of two in **S3** normalizes $p_{ij}$ across unordered node pairs.

Under this model, the probability, $p_{ij}$, of a single co-occurrence event occurring between nodes *i* and *j* by random chance is given by:

$$p_{ij} = \frac{s_i\ s_j}{2(W_{total})^2} \qquad \text{(S3)}$$

To identify the most significant network structures a log-likelihood($\mathcal{L}$) is defined for given subgraph, $E_{sub}$, with total observed weight$W_{sub} = \Sigma_{(i,j)\in E_{sub}} w_{ij}$. $\mathcal{L}$ takes the multinomial form and therefore expresses the probability of the observed configuration under the null; minimizing it isolates the subgraph least probable by chance, i.e. edges whose observations ($w_{ij}$) are most enriched relative to their random probabilities ($p_{ij}$). The objective log-likelihood equation to derive the subgraphs that drives both GLF and SA probabilities is as follows:

$$\mathcal{L}(G_{sub}) = \ln(W_{sub}!) + \sum_{(i,j)\in E_{sub}} \left[w_{ij}\ln(p_{ij}) - \ln(w_{ij}!)\right] \qquad \text{(S4)}$$

We employ GLF and SA independently as stochastic optimization strategies to minimize $\mathcal{L}$. Both chains were initialized by a uniform random draw of edges at the target count (n = 500), which was held fixed thereafter; each proposal removed one member edge and added one non-member edge, both drawn uniformly at random, and was evaluated by the Metropolis criterion on $\Delta\mathcal{L}$. Where the two methods differ is their acceptance probabilities for trading edges: GLF uses a fixed acceptance probability ($T$=1), whereas SA uses a dynamic acceptance probability with scheduled cooling rate, α, (0.999995) with an initial temp (*T*) of 5000. Each search returned the lowest $\mathcal{L}$ configuration encountered rather than its terminal state. GLF and SA were seeded independently (42 and 43) so the chains explore the space without retracing a shared trajectory:

$$P_{GLF}(\text{accept}) = \min(1, e^{-\Delta\mathcal{L}}) \quad \text{(S5)}$$

$$P_{SA}(\text{accept}) = \min(1, e^{-\frac{\Delta\mathcal{L}}{t_k}}) \qquad t_{k+1} = \alpha \cdot t_k \quad \text{(S6),(S7)}$$

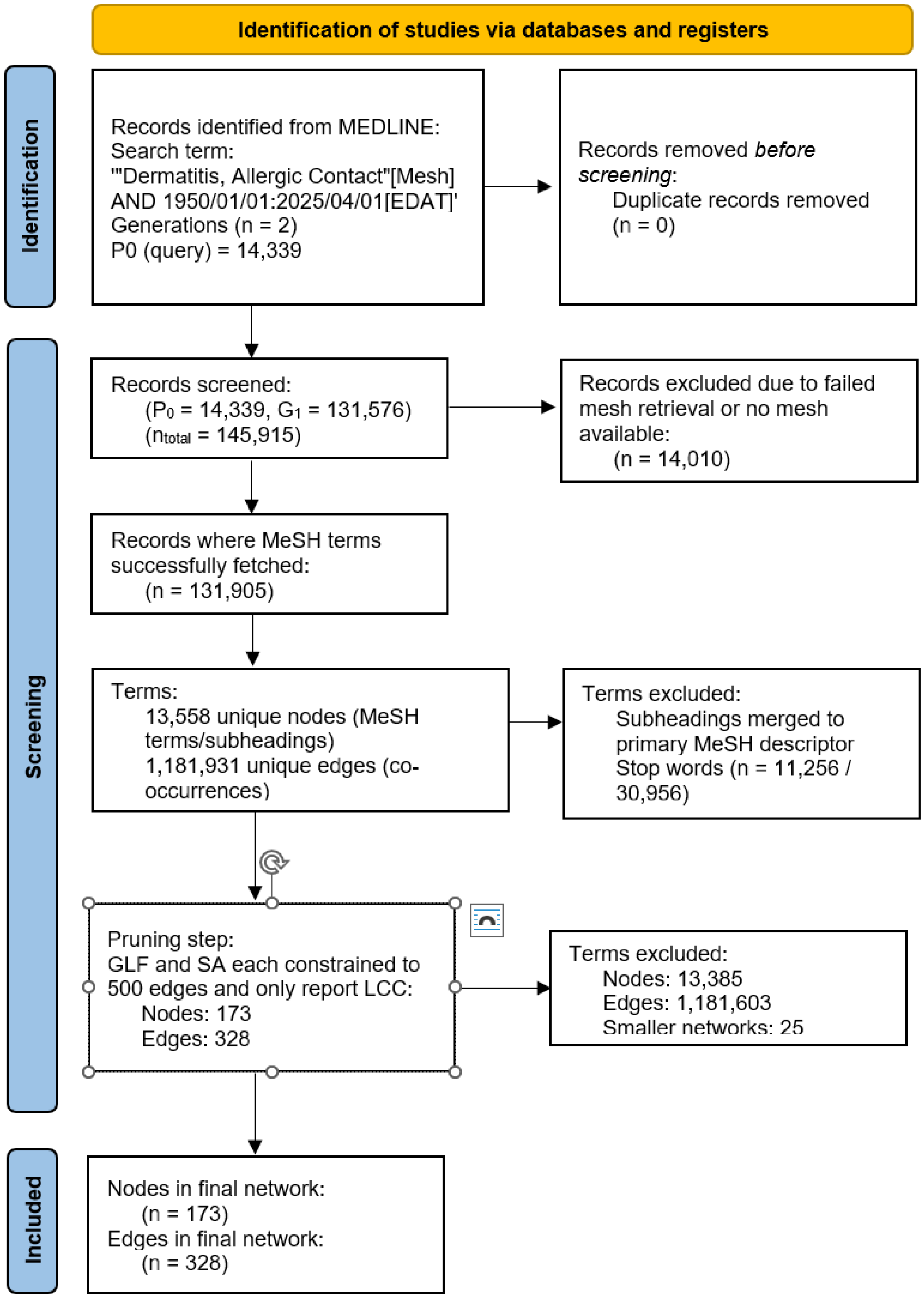


**Supplementary Figure 1:** Adapted PRISMA 2020 flow diagram for machine driven systematic review methodology of NLM's MEDLINE (Pubmed) database. Note that the flow starts from records (publications (PMIDs)) and shifts to MeSH terms during network distillation resulting in MeSH term (nodes) and their co-occurrences (edges).

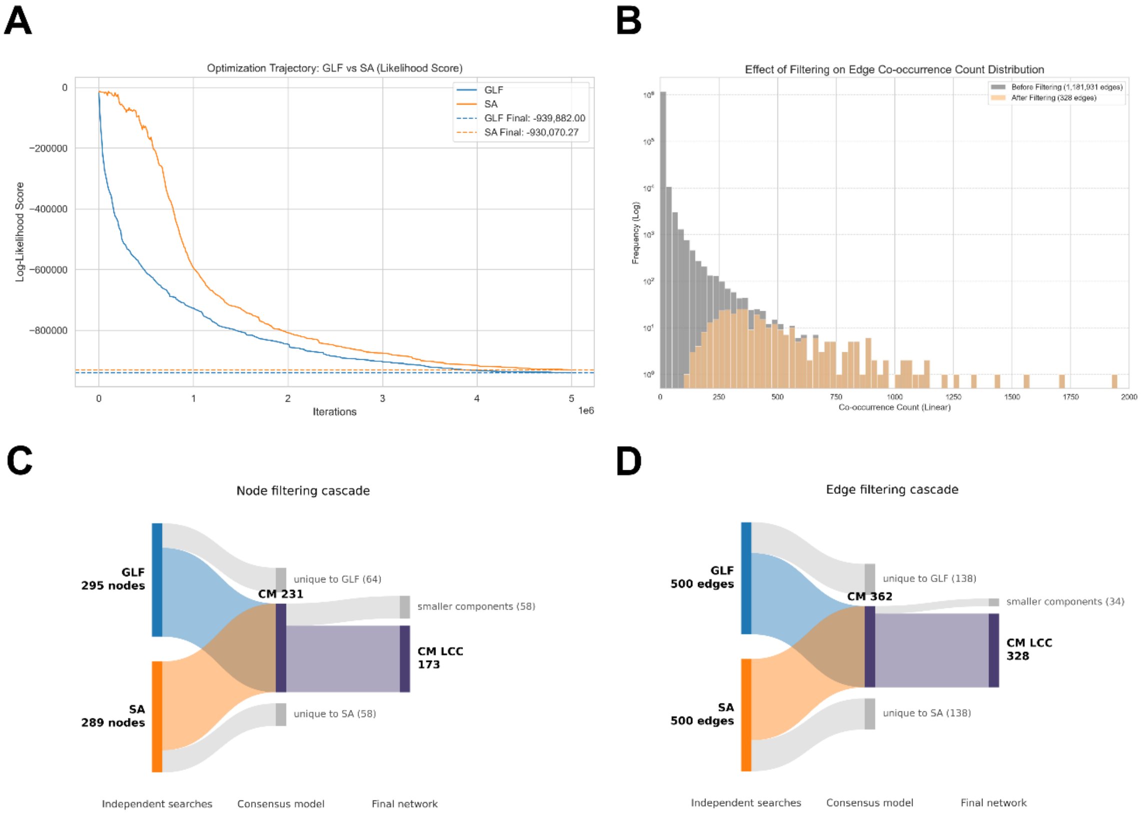


**Supplementary Figure 2:** (**A**) Optimization trajectories of both Global Likelihood Factor (GLF, blue) and Simulated Annealing (SA, orange) over 5x$10^6$ iterations to find an optimal subgraph of edges (*n*=500) representing the log-likelihood of the null model. (**B**) Histogram overlay (log-scale) of MeSH terms co-occurrence count before (grey) and after (orange) consensus filtering. (**C**) Node and (**D**) edge alluvial diagrams of the filtering cascade. Global Likelihood Filter (GLF) and Simulated Annealing (SA) each returned a 500-edge subgraph; the Consensus Model (CM) is the intersection of these two edge sets, from which only the Largest Connected Component (LCC) was retained. Grey flows mark edges and nodes discarded at each stage.

**Supplementary Table 1:** Zero-truncated negative binomial (ZTNB, NB2, intercept-only) regression of the unique MeSH term co-occurrences in the unfiltered Allergic Contact Dermatitis network, fitted by maximum likelihood over all 1,181,931 edges (*statsmodels*, TruncatedLFNegativeBinomialP, truncation = 0). Because the network contains only observed co-occurrences, the support is $Y \geq 1$ and the truncated likelihood accounts for the structurally absent zero class. Overdispersion is decisively confirmed against the zero-truncated Poisson null ($\chi^2$ = 7,448,590, df = 1, $p < 0.001$; variance-to-mean 59.87), motivating the likelihood-based filtering applied downstream. The dispersion parameter is a boundary solution and is not identifiable, $\alpha$ ranged from 900 to 37,529 across five starting values at an identical log-likelihood with a singular Hessian and no valid Wald test, so a no point $\alpha$ is reported. The fitted truncated mean, $E[Y \mid Y \geq 1]$ = 3.0802 co-occurrences per edge (95% CI 3.0557–3.1047; bootstrap 3.059–3.101), is identified and replaces the $\exp(\beta_0)$ of the earlier untruncated model.

```
                   ZTNB (zero-truncated NB2) Regression Results
==============================================================================
Dep. Variable:                  counts   No. Observations:            1181931
Model:                       ZTNB (NB2)   Df Residuals:                1181930
Method:                             MLE   Df Model:                          0
Date:                  Tue, 18 Aug 2026   Pseudo R-squ.:                 0.000
Time:                                 -   Log-Likelihood:         -1.8937e+06
converged:                         True   LL-Null:                -1.8937e+06
Covariance Type:              nonrobust   LLR p-value:                     nan
==============================================================================
                 coef    std err          z      P>|z|      [0.025    0.975]
------------------------------------------------------------------------------
intercept     -8.7400        nan        nan        nan         nan       nan
alpha             nan        nan        nan        nan         nan       nan
==============================================================================

                   Identified quantities and model comparison
==============================================================================
Fitted mean E[Y|Y>=1]:           3.0802   95% CI:                3.0557 - 3.1047
Std error of the mean:           0.0125   Bootstrap (B=200):       3.059 - 3.101
Variance:                        184.41   Variance-to-mean:                59.87
Median / Min / Max:         1 / 1 / 3805   Poisson reference:                1.00
------------------------------------------------------------------------------
ZTNB log-likelihood:         -1,893,678   ZTNB AIC:                    3,787,360
ZTP log-likelihood:          -5,617,973   ZTP AIC:                    11,235,948
LR chi-square (df=1):         7,448,590   LR p-value:                     < 0.001
Delta AIC:                    7,448,588   Delta AIC per edge:               6.30
==============================================================================
```

**Supplementary Table 2:** Sub-network metrics for Global Likelihood Filter (GLF) and Simulated Annealing (SA). Individual density and clustering coefficients via *networkx* Python library constrained to edge$_n$=500. Shared Jaccard and F1 score reported.

| Metric | GLF result | SA result |
|---|---|---|
| Number of Nodes | 295 | 289 |
| Number of Edges | 500 | 500 |
| Network Density | 0.0115 | 0.0120 |
| Average Clustering Coefficient | 0.2791 | 0.2753 |
| Edges Unique to model | 138 | 138 |
| Shared Edges (Consensus) | 362 | 362 |
| Jaccard Index | 0.5674 | 0.5674 |
| F1 Score | 0.7240 | 0.7240 |

**Supplementary Table 3:** Top 5 disconnected components excluding LCC of concensus method. The consensus comprised of 231 nodes and 362 edges distributed across 26 components, of which the LCC (173 nodes) was retained for pathway analysis. Network size and node with highest betweenness reported.

| Rank | Size (# of nodes) | Central theme node | Betweenness | Members |
|---|---|---|---|---|
| #1 | 5 | Terpenes | 0.0008 | Acyclic Monoterpenes; Cyclohexenes; Limonene; Monoterpenes; Terpenes |
| #2 | 4 | Cell Adhesion | 0.0019 | Cell Adhesion; Cell Adhesion Molecules; Endothelium, Vascular; Intercellular Adhesion Molecule-1 |
| #3 | 3 | Pregnancy | 0.0106 | Pregnancy; Pregnancy Complications; Prenatal Exposure Delayed Effects |
| #4 | 3 | Phenols | 0.0007 | Benzhydryl Compounds; Bisphenol A Compounds; Phenols |
| #5 | 3 | COVID-19 | 0.0006 | COVID-19; Coronavirus Infections; Pneumonia, Viral |

**Supplementary Table 4:** Mean Relevancy Score (MRS) rankings of AOP 40-associated MeSH terms within the 173-node consensus Largest Connected Component, ordered by biological stratum and grouped by the key event each term represents. Ranks (1 = highest MRS) are reported for six MRS variants, comprising eigenvector, betweenness and PageRank centrality each computed on the consensus subgraph ("sg") and on the full unfiltered co-occurrence network ("full"). Mean and median rank presented of the twelve terms under each variant, while the top 25 and top 50 rows count how many of the twelve fall within those rank threshold cut-offs. Corpus-wide BEDROC (α = 20) is given for each variant as its corresponding article-retrieval performance.

| Key event | MeSH term | Stratum | eigen (sg) | betw (sg) | PR (sg) | eigen (full) | betw (full) | PR (full) |
|---|---|---|---|---|---|---|---|---|
| Stressor (chemical) | Allergens | Stressor | 7 | 6 | 7 | 41 | 10 | 22 |
| Stressor (chemical) | Irritants | Stressor | 11 | 14 | 16 | 8 | 18 | 6 |
| MIE: covalent protein binding | Haptens | Molecular | 81 | 96 | 90 | 27 | 92 | 51 |
| KE1: keratinocyte activation | Keratinocytes | Cellular | 23 | 30 | 45 | 47 | 13 | 30 |
| KE2: dendritic cell activation | Langerhans Cells | Cellular | 15 | 23 | 25 | 9 | 28 | 10 |
| KE2: dendritic cell activation | Dendritic Cells | Cellular | 25 | 40 | 43 | 55 | 58 | 52 |
| KE3: T-cell activation | T-Lymphocytes | Cellular | 52 | 33 | 61 | 97 | 54 | 89 |
| KE3: T-cell activation | Th1 Cells | Cellular | 20 | 44 | 34 | 22 | 49 | 28 |
| KE3: T-cell activation | Th2 Cells | Cellular | 10 | 21 | 23 | 13 | 38 | 13 |
| AO: allergic contact dermatitis | Dermatitis, Allergic Contact | Adverse Outcome | 4 | 1 | 1 | 4 | 1 | 2 |
| AO: allergic contact dermatitis | Dermatitis, Contact | Adverse Outcome | 6 | 5 | 4 | 10 | 2 | 3 |
| AO: allergic contact dermatitis | Dermatitis, Irritant | Adverse Outcome | 1 | 4 | 2 | 2 | 6 | 1 |

| | | | | | | |
|---|---|---|---|---|---|---|
| *mean rank* | 21.2 | 26.4 | 29.2 | 27.9 | 30.8 | 25.6 |
| *median rank* | 13.0 | 22.0 | 24.0 | 17.5 | 23.0 | 17.5 |
| *terms in top 25* | 10 | 7 | 7 | 7 | 6 | 7 |
| *terms in top 50* | 10 | 11 | 10 | 10 | 9 | 9 |
| *corpus-wide BEDROC* | 0.616 | 0.710 | 0.658 | 0.538 | 0.659 | 0.569 |

**Supplementary Table 5:** Alluvian flow results of edges (n=296) of various Adverse Outcome Pathway (AOP) biological strata excluding Uncategorized edges.

| Source | Target | Weight |
|---|---|---|
| Adverse Outcome | Adverse Outcome | 34 |
| Cellular | Adverse Outcome | 5 |
| Cellular | Cellular | 31 |
| Cellular | Organ | 11 |
| Cellular | Tissue | 11 |
| Molecular | Adverse Outcome | 17 |
| Molecular | Cellular | 27 |
| Molecular | Molecular | 36 |
| Molecular | Organ | 4 |
| Molecular | Tissue | 1 |
| Organ | Adverse Outcome | 12 |
| Organ | Organ | 3 |
| Stressor | Adverse Outcome | 42 |
| Stressor | Cellular | 2 |
| Stressor | Molecular | 13 |
| Stressor | Organ | 3 |
| Stressor | Stressor | 19 |
| Stressor | Tissue | 3 |
| Tissue | Adverse Outcome | 13 |
| Tissue | Organ | 8 |
| Tissue | Tissue | 1 |
